# Effects of vitamin D on inflammatory and oxidative stress responses of human bronchial epithelial cells exposed to particulate matter

**DOI:** 10.1101/351791

**Authors:** Paul E Pfeffer, Haw Lu, Elizabeth H Mann, Yin-Huai Chen, Tzer-Ren Ho, David J Cousins, Chris Corrigan, Frank J Kelly, Ian S Mudway, Catherine M Hawrylowicz

**Author notes:** **Corresponding author:** Dr Catherine M Hawrylowicz, MRC and Asthma UK Centre in Allergic Mechanisms of Asthma, 5th Floor Tower Wing, Guy’s Hospital, King’s College London, London SE1 9RT, UK.

## Abstract

**Background:** Particulate matter (PM) pollutant exposure, which induces oxidative stress and inflammation, and vitamin D insufficiency, which compromises immune regulation, are detrimental in asthma.

**Objectives:** Mechanistic cell culture experiments were undertaken to ascertain whether vitamin D abrogates PM-induced inflammatory responses of human bronchial epithelial cells (HBECs) through enhancement of antioxidant pathways.

**Methods:** Transcriptome analysis, PCR and ELISA were undertaken to delineate markers of inflammation and oxidative stress; with comparison of expression in primary HBECs from healthy and asthmatic donors cultured with reference urban PM in the presence/absence of vitamin D.

**Results:** Transcriptome analysis identified over 500 genes significantly perturbed by PM-stimulation, including multiple pro-inflammatory cytokines. Vitamin D altered expression of a subset of these PM-induced genes, including suppressing *IL6*. Addition of vitamin D suppressed PM-stimulated IL-6 production, although to significantly greater extent in healthy versus asthmatic donor cultures. Vitamin D also differentially affected PM-stimulated GM-CSF, with suppression in healthy HBECs and enhancement in asthmatic cultures. Vitamin D increased HBEC expression of the antioxidant pathway gene *G6PD*, increased the ratio of reduced to oxidised glutathione, and in PM-stimulated cultures decreased the formation of 8-isoprostane. Pre-treatment with vitamin D decreased CXCL8 and further decreased IL-6 production in PM-stimulated cultures, an effect abrogated by inhibition of G6PD with DHEA, supporting a role for this pathway in the anti-inflammatory actions of vitamin D.

**Conclusions:** In a study using HBECs from 18 donors, vitamin D enhanced HBEC antioxidant responses and modulated the immune response to PM, suggesting that vitamin D may protect the airways from pathological pollution-induced inflammation.

## Introduction

Asthma is the most common chronic lung disease with globally increasing prevalence, implying the importance of environmental factors in its aetiology [1]. Vitamin D insufficiency/deficiency and ambient air pollution are two major environmental factors that appear to influence the pathogenesis and stability of asthma [2] [3] [4] [5], as well as other respiratory diseases [6] [7]. However there remains debate resulting from the heterogeneity of findings relating to the effects of these environmental factors on airway pathology [5] [8] [9] [10]. For example, European studies have shown heterogeneity between different cities in the magnitude of the effects of pollution on health outcomes such as hospital admissions for respiratory diseases [9] and asthma incidence [11], despite using standardised analyses. Environment-environment interactions are a major possible explanation for inconsistent results between different patient cohorts but have been little studied, particularly at the mechanistic level. In a recent meta-analysis, latitude of study location influenced associations between air pollutants and severe asthma exacerbations, and latitude is also known to affect sunlight-derived vitamin D production, although this association is complicated by other factors such as hours of daily skin exposure to sunlight [5]. In the urban environment Rosser and colleagues have shown that vitamin D insufficient children, but not those vitamin D sufficient, living close to major roads show an elevated risk of severe asthma exacerbations [12], although the mechanisms by which vitamin D may protect against pollution toxicity remain unclear and the interaction likely complex.

A growing body of research highlights the importance of epithelial immunology in asthma [13]. Evidence shows that inhaled ambient particulate matter (PM) adversely affects the bronchial epithelium through various mechanisms including the imposition of oxidative stress, which stimulates redox sensitive signalling pathways and drives the transcription of pro-inflammatory mediators relevant to asthma and other inflammatory lung diseases [14]. There is evidence that in asthma this pro-oxidant/pro-inflammatory action is superimposed on a background of oxidative stress. For example, Teng and colleagues have shown concentrations of H_2_O_2_ are elevated in exhaled breath condensate of asthmatics compared with controls, with concentrations increasing with asthma severity [15]. Mak and colleagues have reported elevated plasma concentrations of 8-isoprostane, a lipid peroxide marker of oxidative stress, during asthma exacerbations [16]. Indeed oxidative stress has been implicated in many of the key pathophysiological features of asthma, including airways hyper-responsiveness and mucus hypersecretion (reviewed by Zuo *et al.* [17], and Li *et al.* [18]).

There is a large volume of research highlighting the anti-inflammatory effects of vitamin D on the adaptive immune system [19] [20], but limited research as to the effects of vitamin D on the human bronchial epithelium. Hansdottir and colleagues have shown vitamin D to decrease production of pro-inflammatory cytokines by virally infected primary human bronchial epithelial cells (HBECs) [21], but the capacity of vitamin D to affect the epithelial response to urban particulate matter, consistent with it acting as a modifier of PM-induced respiratory effects, to our knowledge has never been examined.

In view of this, we set out to examine whether vitamin D could abrogate urban PM-induced pro-inflammatory responses in primary human bronchial epithelial cells. We elected to commence with an unbiased, transcriptomic analysis with the objective of identifying PM-induced pro-inflammatory cytokines that show distinct patterns of alteration by vitamin D *in vitro*. We studied the epithelial response to both active 1α,25-dihydroxyvitamin D3 (1,25(OH)_2_D3) but also to vitamin D in its circulating precursor form as 25-hydroxyvitamin D3 (25(OH)D3). To examine the effects in a broader human context we studied epithelial cells from both patients with diagnosed asthma and those without. We hypothesised that HBECs from asthmatic, as compared with healthy control subjects, would display an enhanced pro-inflammatory cytokine response to ambient PM exposure. We further hypothesised that the epithelial response to PM exposure could be favourably modified by vitamin D through induction of antioxidant defences.

## Materials and Methods

### Materials

NIST SRM1648a Urban Particulate Matter (National Institute of Standards & Technology, USA) was suspended at 500μg/ml in 5% methanol vehicle. SRM1648a is an urban total particulate matter reference material with mean particle diameter 5.85μm that was collected in the USA [22]. NIST SRM1648a in methanol vehicle, hereafter referred to as NIST, was prepared as follows. Appropriate weights of dry powder particulate matter were placed in a 50ml falcon tube to which 5% methanol (HPLC-grade) in Chelex-100 resin-treated water was added. Following re-suspension and sonication on ice at an amplitude of 15 microns for 30 seconds using a Soniprep 150 plus probe sonicator (MES (UK) Ltd, UK), the resultant suspension was then separated into 1ml aliquots with re-suspension during the aliquoting procedure to avoid PM sedimentation. The aliquots were stored at −70°C. Separate aliquots of the 5% methanol vehicle alone were also used as a vehicle control (VC). A NIST concentration of 50 μg/ml in primary HBEC cultures corresponded to a theoretical surface deposition of 11 μg/ cm^2^.

Ultra-high purity 1,25(OH)_2_D3 (1α,25-dihydroxyvitamin D3; Enzo Life Sciences, UK) was aliquoted dissolved in DMSO (Sigma-Aldrich, UK) at 100μM. Ultra-high purity 25(OH)D3 (25-hydroxyvitamin D3; Enzo Life Sciences, UK) was aliquoted dissolved in sterile absolute ethanol at 1mM. Both were prepared in low light conditions and stored at −80°C prior to use. Dehydroepiandrosterone (DHEA; Sigma-Aldrich, UK) was dissolved in absolute ethanol at 100mM. Poly(I:C) (Invivogen, USA) was dissolved in sterile H_2_O at 1 mg/ml. Sulforaphane (Sigma-Aldrich, UK) was dissolved in DMSO at 300μM.

### Primary human bronchial epithelial cell (HBEC) culture

Primary human bronchial epithelial cells (HBECs) were acquired from Lonza, Switzerland, and locally from endobronchial brushings / biopsies obtained at fibreoptic bronchoscopy with written informed consent of volunteers (Guy’s Research Ethics Committee, South London REC Office 3, REC approval number 09/H0804/108, 16/03/2010) (Table 1).

**Table 1.** Characteristics of donors of HBECs cultured in this study.

| Characteristic | Healthy<br>(n=10) | Asthmatic<br>(n=8) |
| --- | --- | --- |
| from Lonza (n) | 2 | 2 |
| from local bronchoscopy (n) | 8 | 6 |
| Gender | 6 male, 4 female | 3 male, 5 female |
| Age<br>Mean in years (Range) | 30.4<br>(19 – 60) | 34.9<br>(24 – 52) |
| % taking ICS * | - | 83% |
| % taking LABA * | - | 50% |
| % taking LTRA * | - | 33% |
| Daily ICS Dose * Mean (Range)<br>Beclomethasone equivalent | - | 350 (0 – 1000)<br>micrograms |
| Post-BD FEV1 % predicted *<br>Mean (Range) | 105% (97% - 113%) | 111% (86% - 144%) |

BD, bronchodilator. ICS, inhaled corticosteroid. LABA, inhaled long-acting beta-2 agonist. LTRA, leukotriene receptor antagonist. \* % data only for donors enrolled locally (Lonza HBEC donors, for whom these data are not available, excluded).

For locally collected samples, non-smoking volunteers were phenotyped as atopic or non-atopic based upon skin-prick test results to a standard panel of aeroallergens, and as healthy or asthmatic, the latter based on clinical history and confirmation by lung function testing (documented variability in PEF/FEV_1_ of 12% or more in past year, or positive metacholine / mannitol challenge if diagnosis uncertain).

Cells cultures were incubated in flasks in Bronchial Epithelial Cell Growth Medium (BEGM) – constituted by Bronchial Epithelial Basal Medium (BEBM; Lonza, Switzerland) with SingleQuot Supplements (Lonza, Switzerland) of Bovine Pituitary Extract, Insulin, Hydrocortisone, Gentamicin and Amphotericin-B, Retinoic Acid, Transferrin, Triiodothyronine, Epinephrine, and human Epidermal Growth Factor. In later experiments 1% Penicillin-Streptomycin solution (Sigma-Aldrich, UK) and 1% Nystatin suspension (Sigma-Aldrich, UK) were added to passage 0 and passage 1 cultures. Medium was changed every 2 to 3 days.

For samples grown from brushings: at bronchoscopy multiple brushings with an endobronchial brush were made to collect epithelial cells from 10 areas of the bronchial mucosa of normal appearance. The detached cells were washed with warmed BEGM and re-suspended in flasks of warmed BEGM then incubated at 37°C with 5% CO_2_. The medium was changed on the next day and the passage 0 cultures were then subcultured as described.

For samples grown from biopsies: small fragments of endobronchial biopsy with any visible smooth muscle removed by micro-dissection were placed in a universal tube containing 6ml warmed BEGM cell culture medium. Samples were treated with Liberase TL Research Grade (Roche, USA) collagenase (final concentration 62.5 μg/ml) for one hour at 37°C then centrifuged twice in BEGM medium to wash off collagenase before transfer to a cell culture flask and incubation at 37°C with 5% CO_2_. The medium was changed on the next day and the passage 0 cultures were then subcultured as described.

Cell cultures once near-confluent were passaged by detachment of cells using the recommended Trypsin Subculture ReagentPack (Lonza, Switzerland), centrifugation and then re-suspension in fresh BEGM in flasks for further passage or flat-bottomed cell culture plates for experiments.

Flasks and culture plates for primary HBECs obtained locally at bronchoscopy were collagen coated before use – collagen solution was added to each flask / well base and incubated for 2 hours before washing 3 times with sterile H_2_O. Collagen solution comprised of 20μl type 1 calf skin collagen reagent (Sigma-Aldrich, UK) per ml 0.02M acetic acid in sterile H_2_O.

For experiments, once cultures were near-confluent medium was changed to BEGM containing all SingleQuot Supplements except Bovine Pituitary Extract, Retinoic Acid and in later experiments also excluding Hydrocortisone. Bovine Pituitary Extract was removed as it has been found to contain proteins that can provide exogenous protection against oxidative stress [23]. Bovine Pituitary Extract can also contain low concentrations of vitamin D [24]. Retinoic acid was removed as there is some evidence that it may compete with / antagonise vitamin D [25]. For experiments examining effect of vitamin D pre-treatment, 25(OH)D3 was added at 100nM final concentration with/without 100μM DHEA to appropriate wells. After a further 24 hours, cell cultures were stimulated using fresh BEGM (excluding relevant SingleQuot Supplements as above) with stimulation (50μg/ml NIST, 1μg/ml Poly(I:C) with/without 100nM 1,25(OH)D_2_3, 100nM 25(OH)D3 and/or 100μM DHEA) as appropriate. Culture supernatants and lysed cell monolayers were harvested or other assays conducted 4 hours or 24 hours after stimulation of cell cultures. Cell culture experiments were conducted with triplicate wells for each condition in each experiment to allow for variation in plating density with primary cells. For experiments epithelial cells were used between passage 3 and passage 5.

### Gene transcription microarray

Total RNA was extracted from HBECs cultured for 24 hours stimulated with 50 μg/ml NIST and unstimulated, in the presence and absence of 100nM 1,25(OH)_2_D3, from four different donors (two healthy and two asthmatic). Cell culture monolayers were lysed with Qiazol reagent (QIAGEN, USA) then homogenised with QIAshredder columns before storage at −80°C pending extraction of total RNA using a miRNeasy Mini Kit (QIAGEN, USA) according to an adapted manufacturer’s protocol with an off-column DNA digest with TurboDNase (Ambion, USA). mRNA was quantified using a Qubit Fluorimeter (Invitrogen, USA) and quality controlled using an Agilent 2100 Bioanalyser (Agilent Technologies, USA). Samples were prepared for array analysis using SuperScript III Reverse Transcriptase (Invitrogen, USA) and TargetAmp Nano-g Biotin-aRNA Labelling Kit (Epicentre, Illumina, USA). The microarray was conducted on Illumina HT-12v4 Expression BeadChips (Illumina, USA) on an iScan platform (Illumina, USA). Raw array signal intensities were processed in GenomeStudio (Illumina, USA) with a quantile normalisation before export for analysis in GenomicsSuite (Partek, USA). ANOVA analysis was conducted of the normalised data in GenomicsSuite (Partek, USA).

### Real-time quantitative Polymerase Chain Reaction (qPCR) for measurement of gene mRNA expression

Total RNA was extracted as above and quantified using a NanoDrop ND-1000 Spectrophotometer (Thermo Scientific, USA) and then reverse transcribed to cDNA using RevertAid Reverse Transcriptase and complementary reagents (Fermentas, Thermo Scientific, USA). Relative Quantification (RQ) of target genes relative to *18S* rRNA house keeping gene was conducted in triplicates by real-time quantitative polymerase chain reaction (qPCR) using Taqman Universal PCR MasterMix (Applied Biosystems, USA), and an Applied Biosystems Viia 7 real-time thermal cycler. Results were analysed using Viia 7 software (Applied Biosystems, USA). Taqman primers were purchased from Applied Biosystems, USA (S1 Table). Relative expression of mRNA was corrected for efficiency of amplification using the Pfaffl method [26]. In experiments comparing different culture stimulations expression of genes was analysed relative to both *18S* and the unstimulated control cultures (RQ_Unstimulated_).

### Cytokine protein measurement

Culture supernatants from individual wells were stored at −20°C pending measurement of secreted proteins from individual samples. Concentrations of cytokines were measured in culture supernatants by Cytometric Bead Array (CBA; BD Biosciences, USA). Supernatants were incubated for 3 hours with CBA capture beads before beads were washed and then incubated for 2 hours with CBA detection reagent followed by bead sample analysis as per manufacturer’s instructions. Cytokine levels were measured separately for each of the triplicate wells in primary HBEC cultures and then mean averaged before further analysis.

### Oxidative stress assays

8-Isoprostane ELISA: Cell culture supernatant, pooled from triplicate wells, was collected into tubes containing desferrioxamine (DFO) and butylated hydroxytoluene (BHT) and stored at −80°C pending assay. 8-isoprostane was assayed using a commercial ELISA kit (Cayman Chemical, Michigan, USA) according to the manufacturers protocol.

Glutathione Assay: Intracellular total and oxidized glutathione were assessed using the GSH-GSSG-Glo Assay Kit (Promega, Wisconsin, USA) using an adapted protocol. The protocol was adapted as the un-adapted protocol resulted in readings outside the range of the luminometer standard curve. Near-confluent HBECs were cultured for 24 hours with 100nM 25(OH)D3, with 6 wells per condition per experiment, in phenol-red free Airway Epithelial Cell Medium (Promocell, Germany) with all supplements except bovine pituitary extract, retinoic acid and hydrocortisone. After removal of supernatant, one triplicate of wells for each condition was lysed with kit passive lysis buffer (for assay of total glutathione) and the second triplicate for each condition was lysed with kit lysis buffer containing N-ethylmaleimide (for assay of oxidized glutathione). The 50μl lysate in each well was then diluted by addition of 150μl dH_2_O, mixed on an orbital shaker, before 40μl of diluted lysate was transferred to white-sided (flat transparent-bottom) 96 well assay plates. A glutathione standard curve in lysis buffer was prepared according to kit protocol and 40μl added to appropriate wells. 10μl of prepared Luciferin-NT reagent was then added to each well and the plate incubated on an orbital shaker for 5 minutes. The assay was then completed according to the manufacturer’s protocol. A solution of glutathione-S-transferase, dithiothreitol (DTT) and buffer was added to each well and the plate incubated for 30 minutes. Finally, a luciferase containing reagent solution was then added to each well and the plate incubated for 20 minutes prior to reading well luminescence with a GloMax plate luminometer (Promega, Wisconsin, USA). For each condition in each experiment, the concentration of reduced glutathione was calculated by subtracting the mean concentration in the oxidized glutathione triplicate from the mean concentration in the total glutathione triplicate.

### Statistical Analysis

GraphPad Prism 6.0 (GraphPad Software, USA) was used for all statistical tests except for the microarray data. Unless otherwise stated in figure legends, data are presented as mean +/− standard deviation (SD) for normally-distributed data and as box-and-whisker plots showing median, inter-quartile and absolute ranges for non-parametric data. Unless otherwise stated, non-parametric data were analysed using Friedman Tests with Dunn’s multiple-comparisons test and parametric data by repeated-measures ANOVA with Bonferroni corrected post-tests. Significant results shown in figures as follows: *, p ≤ 0.05; **, p ≤ 0.01; ***, p ≤ 0.001. In some experiments results from HBEC cultures from healthy (both atopic and non-atopic) and asthmatic subjects were combined to better analyse the effects of PM and vitamin D across a larger and broader population of individuals.

## Results

### Effects of vitamin D on expression of pro-inflammatory cytokine genes by HBECs exposed to particulate matter

In initial studies to investigate vitamin D mediated inhibition of the pro-inflammatory effects of particulate matter, a gene transcription microarray was conducted on HBECs stimulated with PM for 24 hours, using cultures from four adult donors. Expression of 510 genes was altered (fold-change ≥ ± 1.4) upon stimulation with 50 μg/ml NIST SRM1648a, a reference preparation of urban PM, hereafter referred to as NIST (full microarray data available in S1 Data Appendix). This concentration of NIST had previously been ascertained to induce significantly increased production of pro-inflammatory cytokines after 24 hour cell culture. Of these genes, 49 also showed evidence of regulation (fold-change ≥ ± 1.4) in the presence of the active form of vitamin D, 1,25(OH)_2_D3, at a concentration of 100nM (Fig 1A, S2 Table). Expression of the pro-inflammatory cytokine genes *IL6, CXCL10* and *IL24* was induced by NIST, but suppressed by the addition of vitamin D (Fig 1B). Expression of other pro-inflammatory cytokine genes, for example *IL8*, was induced by NIST, but not altered by addition of vitamin D. In contrast, NIST suppressed and vitamin D increased the expression of *TGFB2*. As previously reported [27], vitamin D induced expression of *IL1RL1* but interestingly this was also induced by NIST stimulation.

**Fig 1.**
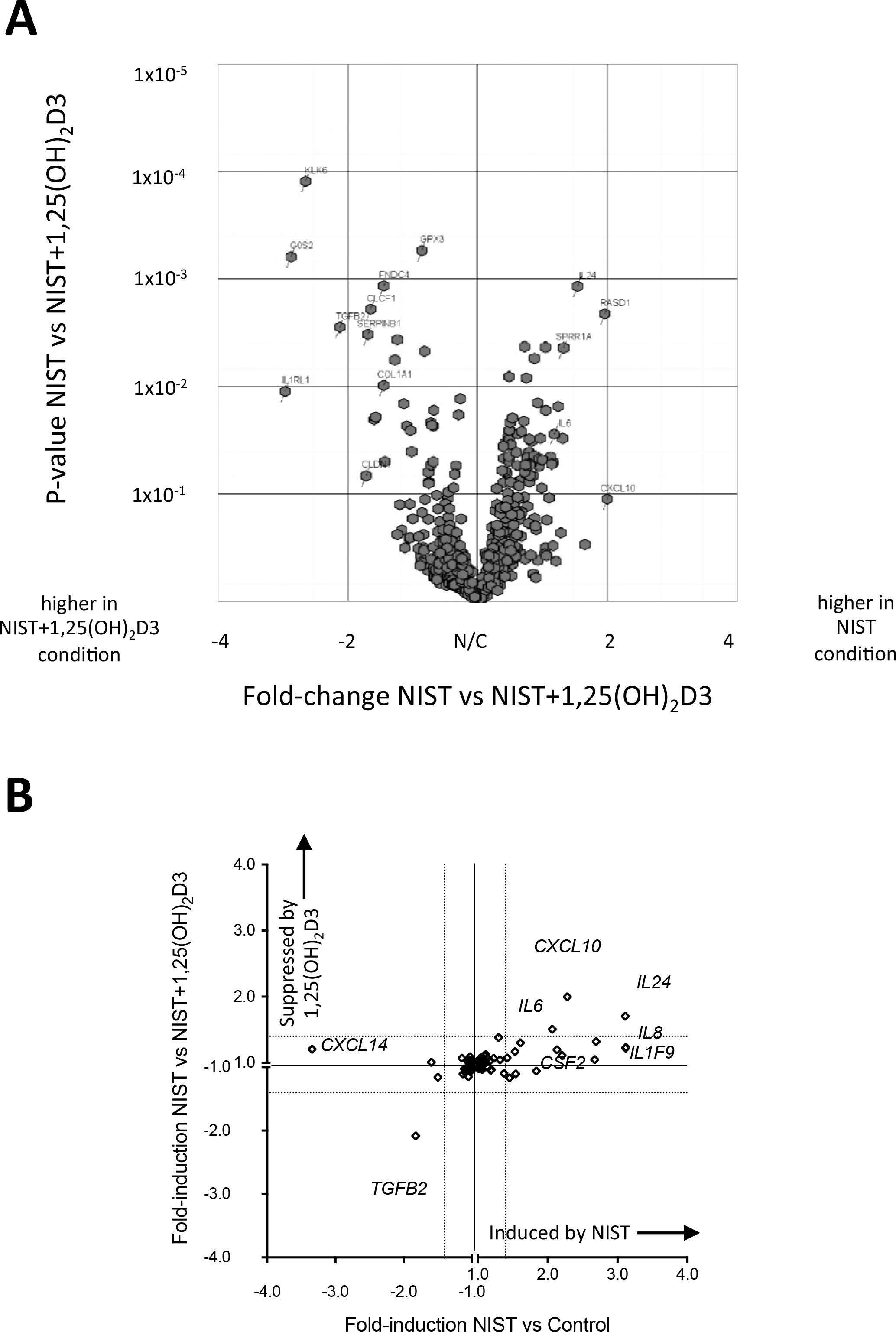
Transcription microarray of 24 hour primary human bronchial epithelial cell (HBEC) cultures stimulated with NIST in the presence/absence of vitamin D. (A) Volcano plot of the 510 genes with ≥ 1.4 fold differential expression comparing 50μg/ml NIST stimulated to unstimulated 24 hour cultures (n=4), showing fold-change in gene expression in NIST stimulated cultures in the presence vs absence of 100nM 1,25(OH)_2_D3 (horizontal axis) plotted again probability of statistical significance for that fold-change (vertical axis). (B) Plot showing microarray results for fold-change in expression of all cytokine genes upon stimulation of HBECs with 50μg/ml NIST (horizontal axis) and fold-change in gene expression upon addition of 100nM 1,25(OH)_2_D3 to NIST-stimulated cultures (vertical axis).

The microarray findings were validated by quantitative real-time PCR using HBEC cultures from a larger number of donors with expression of cytokine genes of interest studied in both 4 hour and 24 hour cultures (Fig 2). Exposure of HBECs to NIST *in vitro* significantly upregulated expression of mRNA encoding *IL6*, *IL8*, *IL24* and *CXCL10* after 4 hours and 24 hours of culture. Expression of *CSF2* was significantly increased by NIST stimulation at 24 hours but not significantly at 4 hours. Co-culture with 100 nM 1,25(OH)_2_D3 reduced *IL6*, *IL24* and *CXCL10* expression at both time points although the effect of vitamin D on *CXCL10* expression at 24 hours did not attain statistical significance. NIST-induced *IL8* expression was not significantly altered by vitamin D at 4 hours but was decreased by 1,25(OH)_2_D3 at 24 hours.

**Fig 2.**
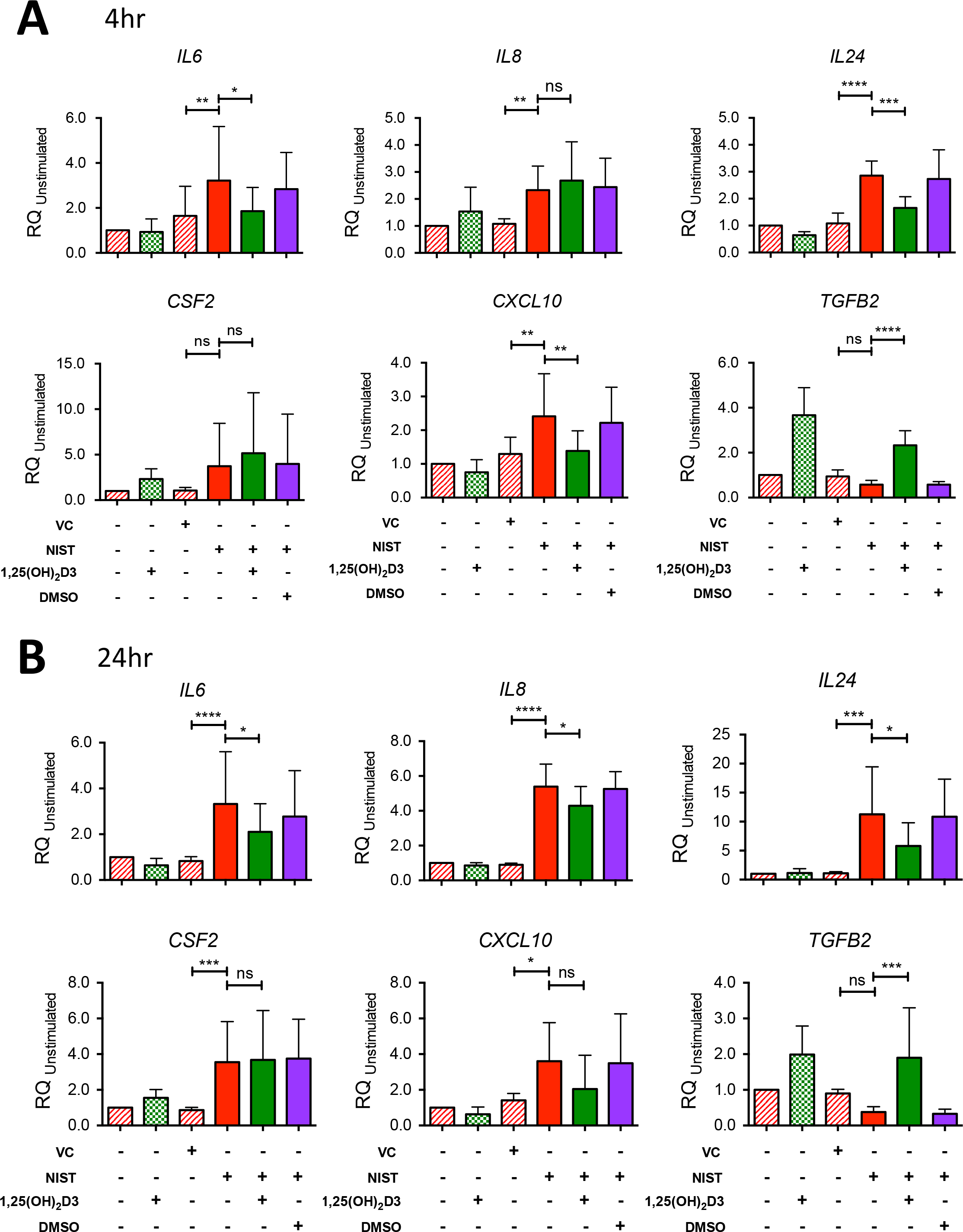
Confirmation by qPCR of the effect of NIST and vitamin D on expression of cytokine genes in HBEC cultures. (A) Gene mRNA expression at 4 hours relative to *18S* and the unstimulated Control condition in HBEC cultures stimulated with 50μg/ml NIST in the presence/absence of 100nM 1,25(OH)_2_D3. *IL6*, n=7; *IL8*, n=7; *IL24*, n=6; *CFS2*, n=8; *CXCL10*, n=6; *TGFB2*, n=6. (B) mRNA expression at 24 hours. *IL6*, n=9; *IL8*, n=7; *IL24*, n=6; *CFS2*, n=8; *CXCL10*, n=6 (one outlying replicate excluded); *TGFB2*, n=6. All repeated-measures ANOVAs and post-tests with Bonferroni corrections as shown. VC; vehicle control for NIST. Statistical significance as follows: *, p ≤ 0.05; **, p ≤ 0.01; ***, p ≤ 0.001; ****, p ≤ 0.0001.

### Comparison of the effects of vitamin D on PM-stimulated epithelial cytokine production by HBECs from healthy and asthmatic individuals *in vitro*

Production by HBECs of the pro-inflammatory cytokines IL-6, CXCL8 and GM-CSF (encoded by the genes *IL6*, *IL8* and *CSF2* respectively) was chosen for further scrutiny given that all three mediators are featured in asthma pathophysiology and their expression was induced by NIST-stimulation, but differently affected by the additional presence of vitamin D. Culture of primary HBECs from 14-17 donors per experiment (approximately equal numbers of healthy and asthmatic subjects) for 24 hours in the presence of 50 μg/ml NIST significantly increased the supernatant concentrations of IL-6, CXCL8 and GM-CSF as compared with vehicle control, consistent with the gene expression data (Fig 3A). In the additional presence of 100nM 1,25(OH)_2_D3 throughout the 24 hour stimulation, production of IL-6 was significantly inhibited, but that of CXCL8 or GM-CSF not significantly altered (Fig 3A).

**Fig 3.**
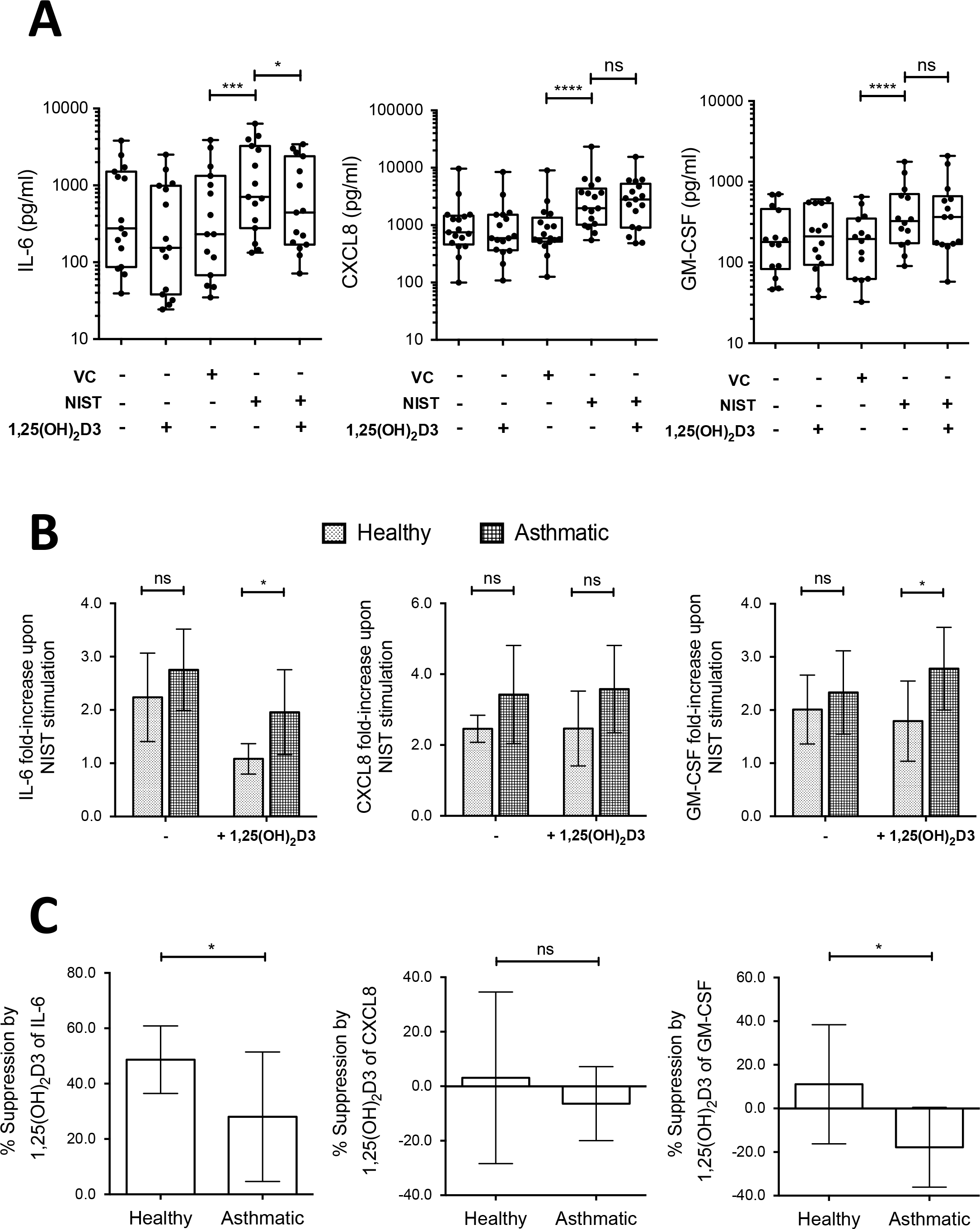
Effects of 1,25(OH)_2_D3 on production of IL-6, CXCL8 and GM-CSF by NIST stimulated HBEC cultures. (A) Addition of 100nM 1,25(OH)_2_D3 reduced production of IL-6 by primary HBEC cultures stimulated for 24 hours with 50μg/ml NIST, but not CXCL8 or GM-CSF. VC: NIST vehicle control. n=14-17. (B) Fold-increase in production of IL-6, CXCL8, and GM-CSF above that in unstimulated cultures upon stimulation with 50μg/ml NIST in the presence/absence of 1,25(OH)_2_D3, by disease status. Two-way ANOVAs with Bonferroni-corrected post-tests. n=8-9 healthy, n=7-8 asthmatic. (C) Percentage suppression by 1,25(OH)_2_D3 of cytokine production in PM-stimulated cultures of HBECs from healthy donors compared to asthmatic donors. Un-paired t-tests. n=8-9 healthy, n=7-8 asthmatic. Statistical significance as follows: *, p ≤ 0.05; **, p ≤ 0.01; ***, p ≤ 0.001, ****, p ≤ 0.0001.

Spontaneous production of all three of these cytokines by HBECs from both the non-diseased and asthmatic donors varied considerably between individuals, with no significant difference between the asthmatic group and healthy group (S1 Fig). To allow for this inter-individual variation, NIST responses were also examined in terms of fold-changes above spontaneous production in unstimulated cultures. Contrary to our hypothesis, NIST-induced fold-increases in cytokine production were not significantly greater in HBECs from asthmatic compared to healthy subjects in the absence of vitamin D (Fig 3B). Nevertheless, 1,25(OH)_2_D3 exhibited a significantly greater capacity to suppress NIST-stimulated IL-6 production by HBECs from the non-diseased, compared with the asthmatic donors (Fig 3C). There was also a significant difference in the effect of 1,25(OH)_2_D3 on NIST-induced GM-CSF production, with suppression in cell cultures from the non-diseased donors, but enhancement in those from the asthmatic donors. As a result, in NIST-stimulated vitamin D treated cultures induction of IL-6 and GM-CSF production was significantly greater in asthmatic than in healthy HBECs (Fig 3B). There was no significant difference in the effect of vitamin D on CXCL8 production by HBECs from the non-diseased and asthmatic donors.

We examined the possibility that these differences in the effects of 1,25(OH)_2_D3 on IL-6 and GM-CSF protein production reflected differences in early induction of cytokine gene expression. No significant differences were evident between cultures of HBECs from healthy and asthmatic donors in NIST-stimulated induction of *IL6* or *CSF2* at 4 hours and 24 hours, compared in the presence and absence of 1,25(OH)_2_D3 (two-way ANOVA analyses, S2 Fig). Additionally, gene expression was compared with the capacity of vitamin D to suppress cytokine production. A proportion of HBEC donor cultures showed greater induction of *CSF2* at 4 hours with less suppression of GM-CSF production at 24 hours by vitamin D (S2 Fig), and notably these were from asthmatic donors. A similar pattern was not evident for IL-6 (S2 Fig).

### Comparison of capacity for 25(OH)D3 and 1,25(OH)_2_D3 to modulate HBEC cytokine responses

Exposure of HBECs to NIST (50 μg/ml, 24 hours) significantly increased the expression of the gene *CYP27B1* (S3 Fig), encoding the cytochrome P450 enzyme that converts 25(OH)D3 to active 1,25(OH)_2_D3. However, the magnitude of induction of *CYP27B1* with NIST exposure was significantly lower than that observed with the established TLR3 agonist Poly(I:C) (mean (SD) 1.2 (0.22) fold induction with NIST compared to 5.4 (3.3) fold induction with TLR3 agonist) (S3 Fig). Similarly, exposure to NIST modestly, but significantly, increased expression of *VDR*, encoding the vitamin D receptor (S3 Fig). Concordantly, both 25(OH)D3 and 1,25(OH)_2_D3 at 100nM concentrations exerted similar effects on cytokine expression by NIST-stimulated HBECs (S3 Fig).

Given the distinct effects of vitamin D on cytokine gene expression in HBECs from healthy compared to asthmatic donors, we were interested to investigate whether this reflected differences in their capacity to respond to vitamin D. We therefore assessed expression of *CAMP*, the gene encoding cathelicidin, and *CYP24A1*; two genes known to be strongly induced by 1,25(OH)_2_D3 (Fig 4). No significant differences were apparent between cells from healthy and asthmatic donors in the capacity of 100nm 1,25(OH)D_2_3, in the presence of NIST, to induce increased expression of *CYP24A1* or *CAMP* relative to unstimulated controls (Fig 4A,C). Similarly expression of the target genes relative to *18S* in the stimulated condition were not significantly different between cultures from healthy and asthmatic donors (Figs 4B,D).

**Fig 4.**
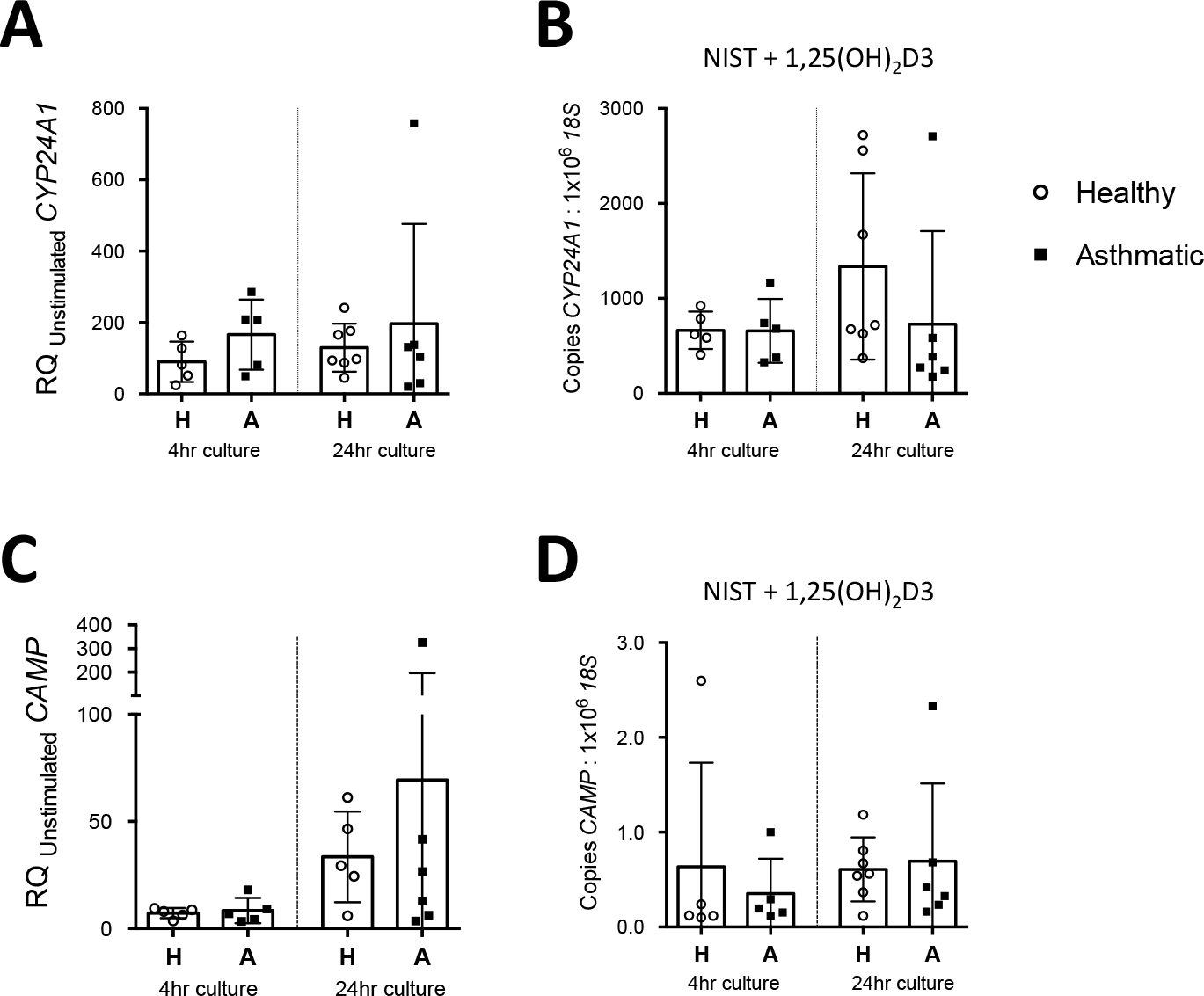
Comparison of expression of vitamin D axis genes between HBECs from healthy and asthmatic donors. (A) Induction of *CYP24A1* by NIST with 100nM 1,25(OH)_2_D3, relative to the unstimulated control condition, in 4 hour and 24 hour HBEC cultures. H, Healthy donor HBECs; A, Asthmatic donor HBEC cultures. 4hr healthy, n=5; 4hr asthmatic, n=5; 24hr healthy, n=7; 24hr asthmatic, n=6. (B) Expression of *CYP24A1* as measured by qPCR relative to *18S* in 4 hour and 24 hour HBEC cultures stimulated with 50μg/ml NIST and 100nM 1,25(OH)_2_D3. 4hr healthy, n=5; 4hr asthmatic, n=5; 24hr healthy, n=7-8; 24hr asthmatic, n=6. (C) Induction of *CAMP* by NIST with 100nM 1,25(OH)_2_D3, relative to the unstimulated control condition, in 4 hour and 24 hour HBEC cultures. 4hr healthy, n=5; 4hr asthmatic, n=5; 24hr healthy, n=5; 24hr asthmatic, n=6. (D) Expression of *CAMP* as measured by qPCR relative to *18S* in 4 hour and 24 hour HBEC cultures stimulated with 50μg/ml NIST and 100nM 1,25(OH)_2_D3. 4hr healthy, n=5; 4hr asthmatic, n=5; 24hr healthy, n=7; 24hr asthmatic, n=6.

### Effects of vitamin D on HBEC oxidative stress and antioxidant responses

We next sought to establish whether the anti-inflammatory effects of vitamin D following PM challenge were related to its upregulation of antioxidant pathways. Culture of HBECs with 100nM 25(OH)D3 for 24 hours significantly increased the intracellular ratio of reduced to oxidised glutathione concentrations (GSH:GSSG; Fig 5A), and in NIST-stimulated cultures 100nM 25(OH)D3 significantly decreased the production of the lipid oxidation product 8-isoprostane (Fig 5B). In further support of an antioxidant action of vitamin D, the antioxidant enzyme-inducing compound sulforaphane similarly suppressed production of IL-6, but not CXCL8 or GM-CSF (Fig 6) in NIST-stimulated cultures.

**Fig 5.**
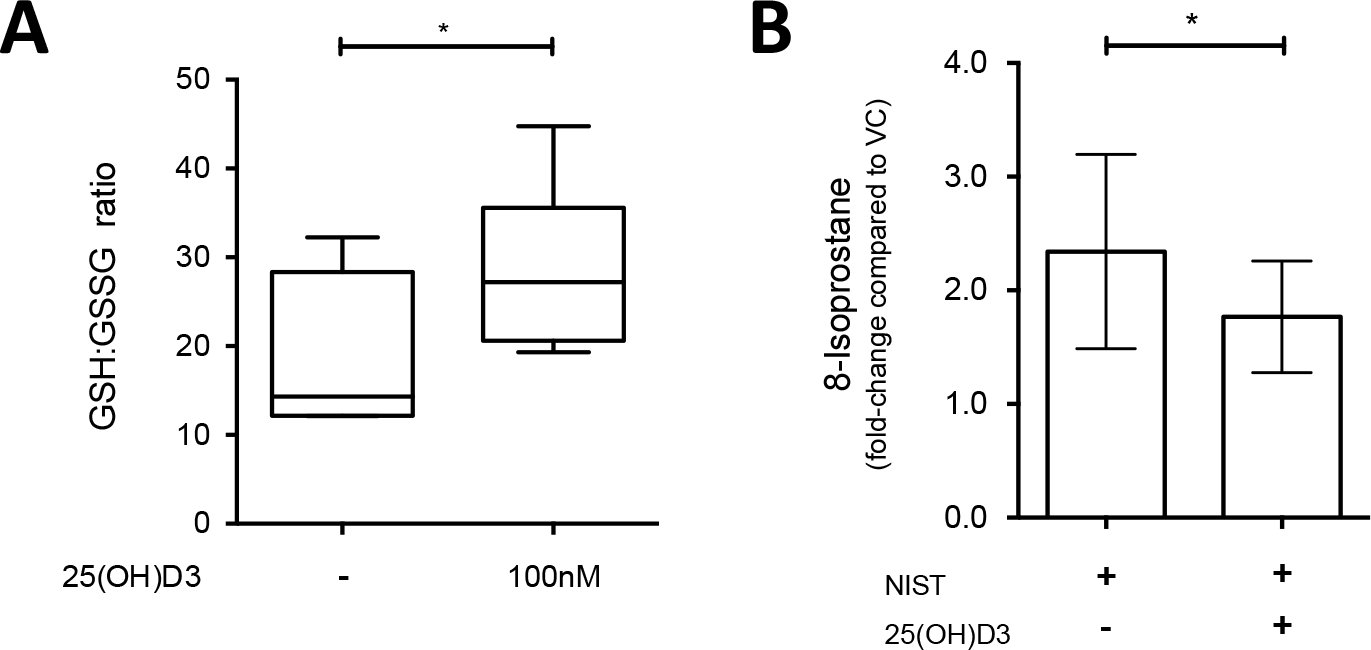
Effect of 25(OH)D3 on antioxidant responses in primary HBEC cultures. (A) Ratio of reduced (GSH) to oxidised (GSSG) glutathione in 24 hour cultures of HBECs treated with 100nM 25(OH)D3; n=6. (B) Fold-increase in 8-isoprostane levels in culture supernatants from primary HBECs cultured with 50μg/ml NIST with/without 100nM 25(OH)D3 for 24 hours, compared to VC control cultures; ratio paired t-test, n=8. Statistical significance as follows: *, p ≤ 0.05.

**Fig 6.**
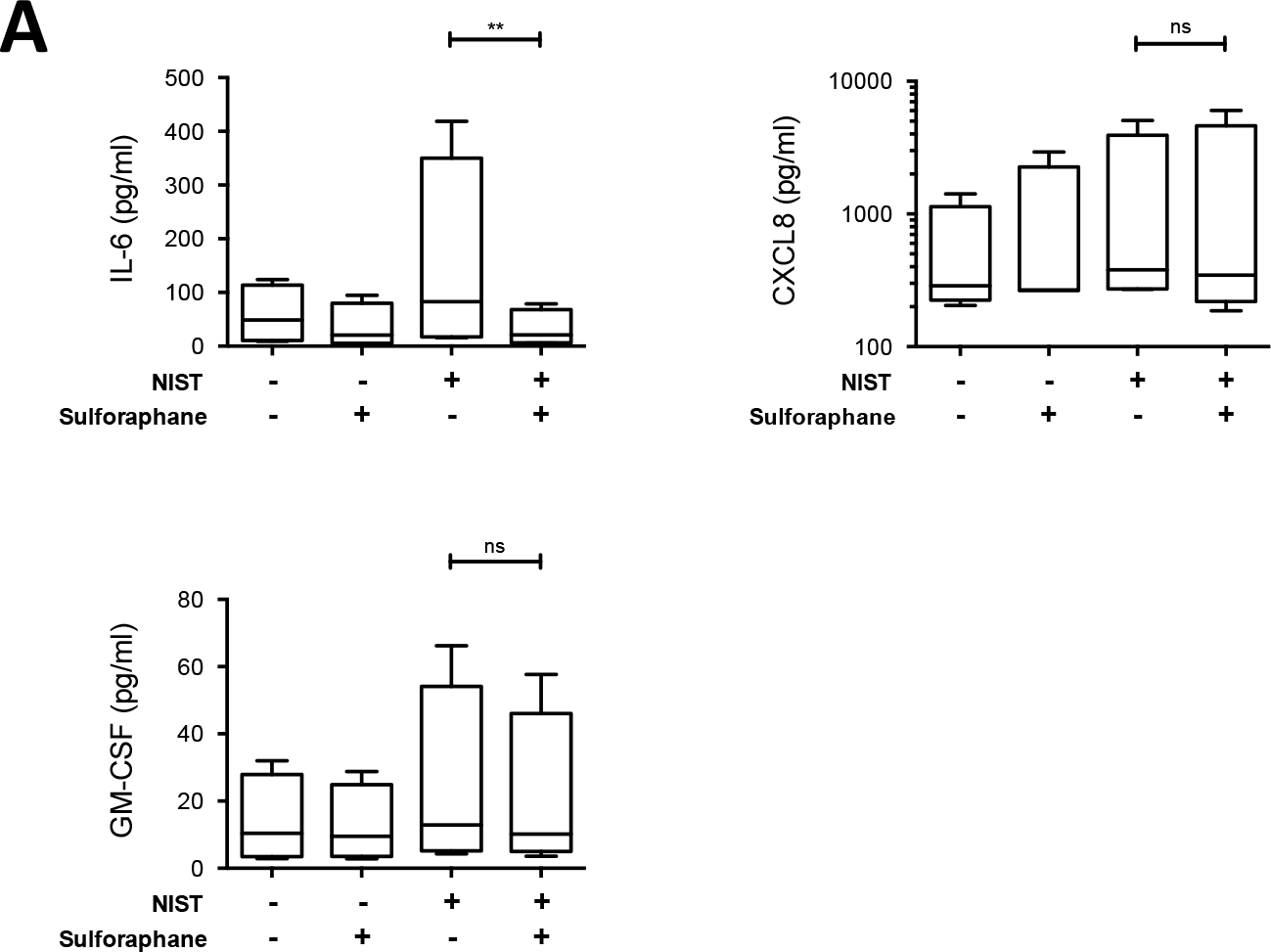
Effect of antioxidant enzyme-inducing sulforaphane on production of IL-6, CXCL8 and GM-CSF by NIST-stimulation HBECs. Cytokines produced by primary HBECs in submerged cultures for 24 hours stimulated with NIST at 50μg/ml and/or sulforaphane at 3μM. Friedman’s tests with Dunn’s multiple comparisons tests; n=4. Statistical significance as follows: **, p ≤ 0.01.

The transcriptomic analysis described above revealed that expression of the gene *G6PD*, which encodes the key antioxidant enzyme glucose-6-phosphate dehydrogenase (G6PD) [28], was significantly upregulated by vitamin D (1.82 fold-increase for NIST + 1,25(OH)_2_D3 compared to NIST alone; Fig 1). The capacity of 1,25(OH)_2_D3 to elevate expression of *G6PD* was confirmed by qPCR in HBEC cultures treated with 100 nM 1,25(OH)_2_D3 for 4 hours and 24 hours (both p<0.001; Fig 7A and B). 25(OH)D3 similarly enhanced *G6PD* expression in a concentration-dependent manner (Fig 7B). There were no significant differences between asthmatic and healthy donor cultures in spontaneous expression of *G6PD* in unstimulated cultures at 4 hours and 24 hours (Fig 7C). Additionally there were no significant differences in the capacity of NIST with/without 1,25(OH)_2_D3 to induce expression of *G6PD* above that in unstimulated cultures (Fig 7D) in HBECs from asthmatics compared to healthy donors.

**Fig 7:**
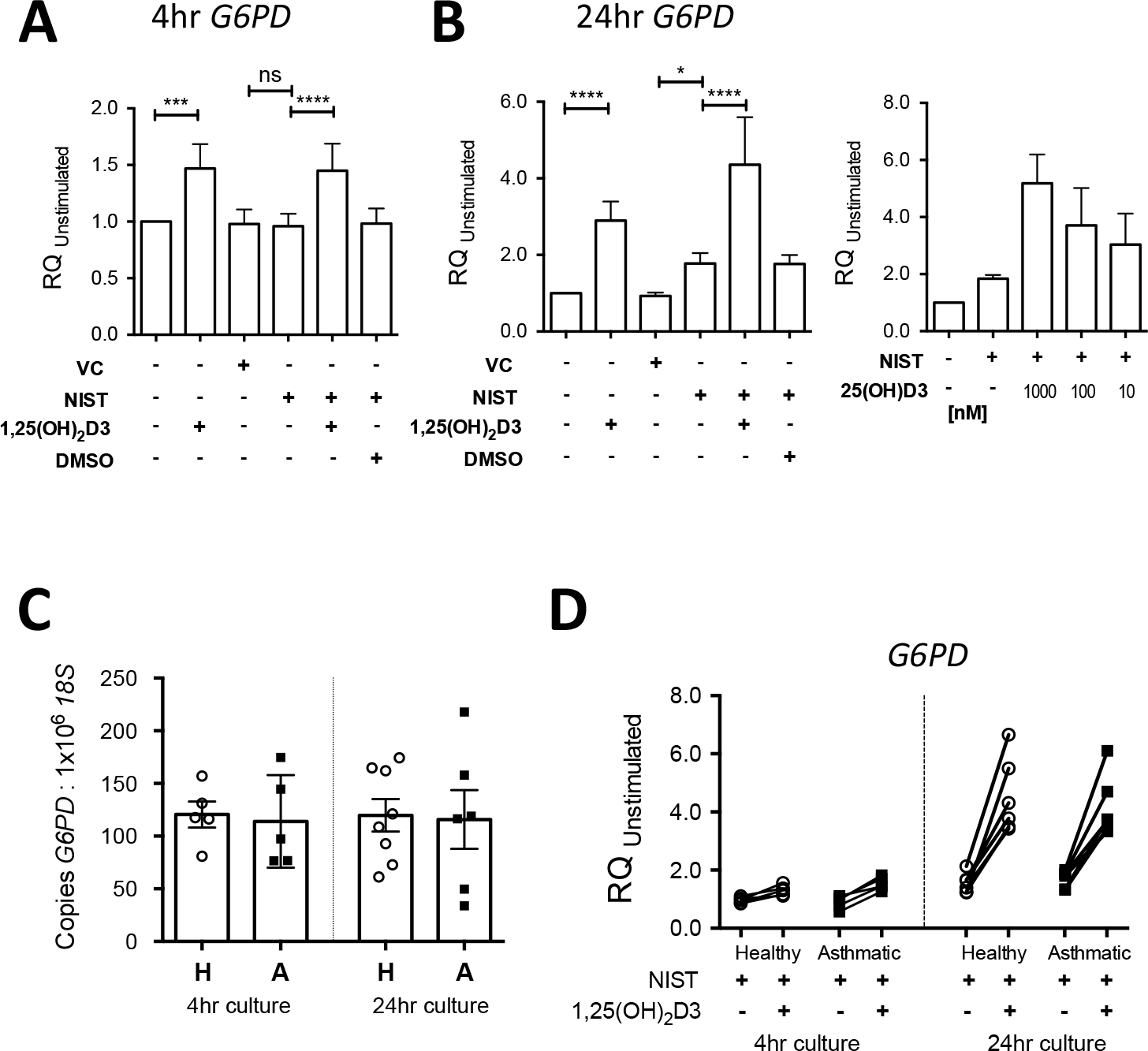
Capacity of vitamin D to enhance primary HBEC expression of *G6PD*. (A) Expression of *G6PD* in 4 hour cultures of primary HBECs stimulated with 50μg/ml NIST with/without 100nM 1,25(OH)_2_D3 (n=6). (B) Expression of *G6PD* in 24 hour cultures of primary HBECs stimulated with 50ug/ml NIST with/without 100nM 1,25(OH)_2_D3 (n=6) or a concentration series of 25(OH)D3, n=5. (C) Expression of *G6PD* as measured by qPCR relative to *18S* in unstimulated 4 hour and 24 hour HBEC cultures. H, Healthy donor HBECs; A, Asthmatic donor HBEC cultures. 4hr healthy, n=5; 4hr asthmatic, n=5; 24hr healthy, n=8; 24hr asthmatic, n=6. (D) Induction of *G6PD* by 50μg/ml NIST in the presence / absence of 100nM 1,25(OH)_2_D3 in 4 hour and 24 hour cultures of HBECs from healthy and asthmatic donors. 4hr healthy, n=5; 4hr asthmatic, n=5; 24hr healthy, n=6; 24hr asthmatic, n=6. Statistical significance as follows: *, p ≤ 0.05; **, p ≤ 0.01; ***, p ≤ 0.001; ****, p ≤ 0.0001.

### Assessing the capacity of vitamin D pre-treatment to suppress pro-inflammatory cytokine production by NIST-stimulated HBECs

Since greater induction by vitamin D of *G6PD* was evident in the HBECs at 24 hours compared with 4 hours in the preceding experiments, we finally investigated whether 24 hours pre-treatment of HBEC cultures with vitamin D prior to particulate challenge would further augment its capacity to decrease production of pro-inflammatory cytokines. Accordingly, we compared the effects of concurrent treatment with 100 nM 25(OH)D3 (as in the previous experiments above) with those of additional treatment of the HBECs with 100 nM 25(OH)D3 for 24 hours prior to exposure to NIST-stimulation (Fig 8A). Additional vitamin D pre-treatment of NIST-stimulated HBECs significantly increased the percentage suppression of IL-6 production (Fig 8A). Vitamin D pre-treatment also suppressed NIST-induced CXCL8 production, a phenomenon not observed when vitamin D was added only concurrently with NIST. However, suppression of IL-6 production (44.78%, 18.45% - 71.10% [mean, 95% confidence interval]) was greater than that of CXCL8 (22.36%, −1.42% −46.14%). In contrast, there was no significant effect on NIST-induced GM-CSF production.

The possible contribution of enhanced expression of G6PD to this phenomenon was assessed using the G6PD inhibitor dehydroepiandrosterone (DHEA). DHEA at a concentration of 100 μM significantly abrogated the ability of 100nM 25(OH)D3, applied for 24 hours in advance and with NIST stimulation, to suppress CXCL8 production in NIST-stimulated HBEC cultures with evidence of a trend towards decreasing suppression of IL-6 (Fig 8B).

**Fig 8:**
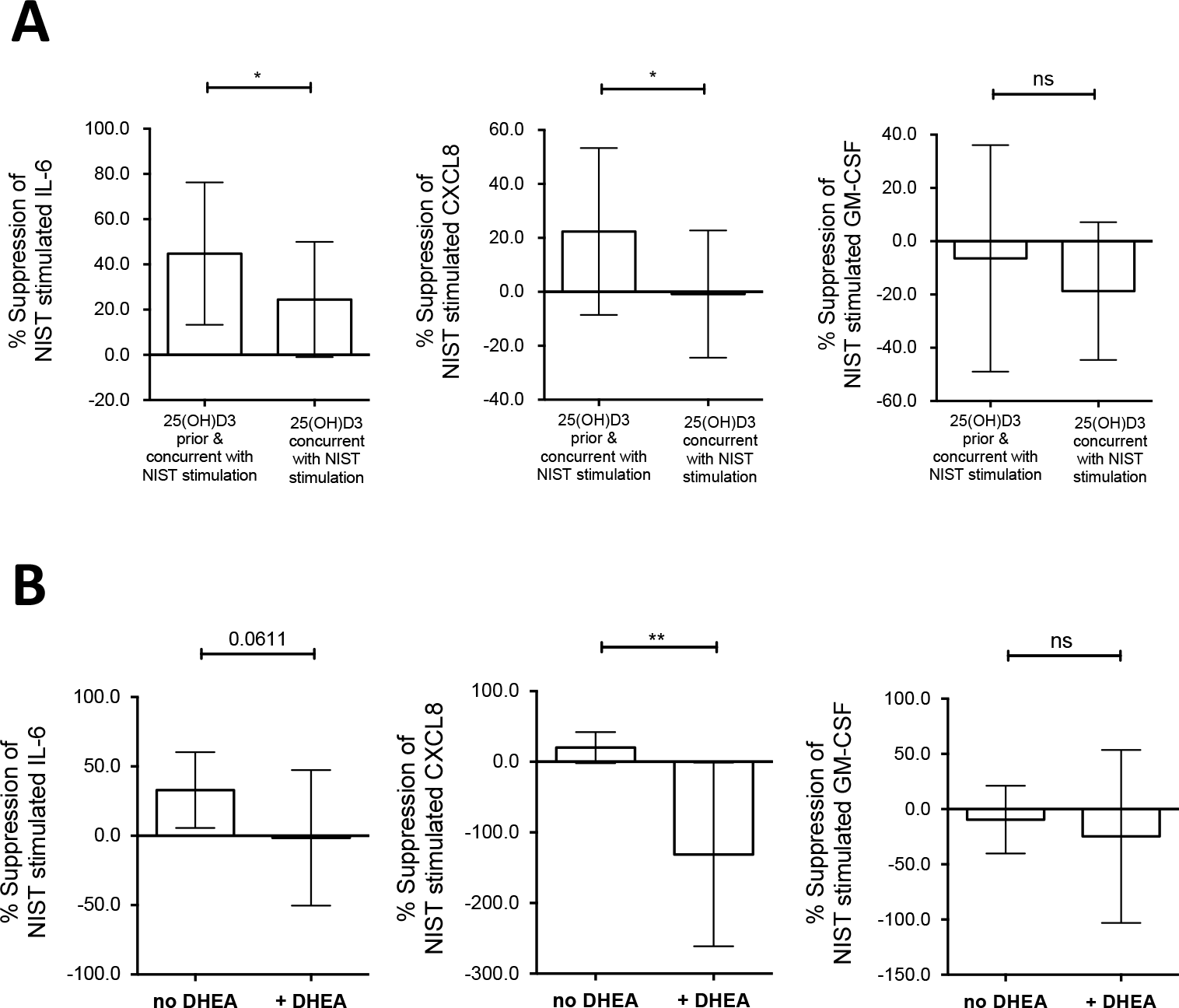
Effect of vitamin D pre-treatment of HBEC cultures on suppression of NIST induced IL-6 and CXCL8. (A) Percentage suppression of cytokine production in 50μg/ml NIST stimulated 24 hour cultures by concurrent and additional 24 hour pre-treatment with 100nM 25(OH)D3 compared to concurrent only; n=7-9 (3-4 Healthy, 4-5 Asthmatic). Two-tailed paired t-tests. (B) Percentage suppression of cytokine production by 100nM 25(OH)D3 concurrent and pre-treatment in 50μg/ml NIST stimulated HBEC cultures, in the absence or presence of the G6PD inhibitor DHEA at 100μM; n=7-9 (2-3 Healthy, 5-6 Asthmatic). Two-tailed paired t-tests. Statistical significance as follows: *, p ≤ 0.05; **, p ≤ 0.01.

## Discussion

Particulate matter air pollution and vitamin D deficiency are environmental factors that have both been implicated in the current ‘epidemic’ of asthma [1] [7] [19]. There is increasing appreciation of the importance of the interaction between multiple environmental factors, defining an individual’s total exposome, in determining health effects, but to date the interactive effect of vitamin D on responses to air pollution has received little attention [5] [29] [30]. In the present study we demonstrate that vitamin D abrogates pro-inflammatory effects of urban PM on primary human bronchial epithelial cells, suppressing production of key PM-stimulated pro-inflammatory cytokines, such as IL-6, through a mechanism at least in part dependent on enhancement of antioxidant pathways by vitamin D. Although epithelial cells from both healthy and asthmatic donors were responsive to vitamin D, the capacity for vitamin D to suppress PM-induced production of pro-inflammatory cytokines was impaired in cells from patients with asthma compared to healthy subjects in this study. A microarray was employed to generate an unbiased depiction of the effects of these environmental factors on epithelial cytokine responses and additionally identified further vitamin D promoted genes such as *G6PD* and *TGFB2*.

Multiple pro-inflammatory cytokines were induced by PM-stimulation, however, the effect of vitamin D on cytokine responses was not uniform with *IL6*, *CXCL10* and *IL24* transcripts suppressed by vitamin D, whereas others such as *IL8* and *CSF2* were not similarly inhibited. We subsequently focused on IL-6, CXCL8 and GM-CSF as these cytokines are well-established to be up-regulated by PM exposure, as well as being implicated in the pathogenesis of asthma, with elevated concentrations seen in lungs of asthmatic individuals [31] [32] [33] [34]. IL-6 is implicated in systemic inflammation [35] [36] [37]. CXCL8 (IL-8) has a major role in neutrophil chemotaxis and activation, with also evidence for a role in eosinophil chemotaxis [38]. GM-CSF is a known HBEC-derived mediator that enhances the pro-inflammatory effects of PM on dendritic cells (DCs) [39] and promotes eosinophil responses [40], but may protect against oxidative injury under a variety of situations [41] [42].

Vitamin D suppressed PM-induced IL-6 production to a greater extent in cultured epithelial cells from healthy as compared with asthmatic donors. In contrast, in cultures from asthmatic donors vitamin D modestly increased GM-CSF production. Why healthy and asthmatic HBECs in this study responded differently to vitamin D is unclear, and future research is needed to address this issue and verify findings in a larger cohort. Nevertheless, this novel finding suggests that future research studying the role of vitamin D in disease pathology should be conducted using primary cells from individuals with those diseases given the important differences apparent between the vitamin D responses of cells from healthy and diseased donors.

Vitamin D exhibits both inhibitory and inductive effects of importance in chronic airway diseases and is therefore likely to influence the response of the lung to pollution exposure. For example, any action of vitamin D to decrease PM-stimulated IL-6 production is likely to be clinically relevant and beneficial. Higher concentrations of IL-6 have been shown in induced sputum from asthmatic patients compared to controls [31], in the plasma of asthmatic patients [43], and a genome-wide association study has previously highlighted the IL-6 receptor gene locus as a possible risk locus for asthma [44]. *In vivo* human exposure to PM has been shown to increase both airway and systemic concentrations of IL-6, with associated acute neutrophilic inflammation [35] [45]. IL-6 can stimulate proliferation of lymphocytes [46], inhibit the action of regulatory T lymphocytes (Tregs) [37] and conversely enhance Th17/IL-17 synthesis [47], which is associated with neutrophilic inflammation and steroid insensitivity in asthma [48]. Elevated IL-17A has been reported with both *in vitro* and *in vivo* particulate matter exposure [49] [50]. Furthermore, the benefit of reducing pollution-induced IL-6 is likely broader than its effect on airway inflammation, since pollution-induced vascular dysfunction and increased blood coagulation have been shown to be IL-6 dependent in murine models [35] [36].

In contrast vitamin D up-regulated expression of *TGFB2*, a cytokine with the capacity to both inhibit inflammation and promote wound healing/fibrosis, with PM having the reciprocal effect. Whilst PM-stimulation creates an immune microenvironment that promotes pro-inflammatory leukocyte responses, vitamin D appears to have the capacity to regulate the production by epithelial cells of multiple mediators that act on the adaptive immune system, such as the cytokines above and as we recently described sST2 [27], to produce a less inflammatory microenvironment.

Vitamin D enhanced expression of the gene *G6PD* encoding glucose-6-phosphate dehydrogenase, the rate-limiting step in the generation of NADPH, necessary for the action of critical antioxidant enzymes including those involved in the production of reduced glutathione [28]. In keeping with this action to enhance antioxidant pathways, vitamin D increased the ratio of intracellular reduced to oxidized glutathione and decreased the production of 8-isoprostane. This action of vitamin D is consistent with previous evidence that it can protect various other types of epithelial cell from oxidative stress *in vitro*, including H_2_O_2_-treated prostatic and breast epithelial cells [51] [52] and CoCl_2_-treated trophoblasts [53]. Similarly, using immortalised epithelial cell lines, notwithstanding that immortalised cell lines are known to manifest distinctly different responses to vitamin D [54], it has been shown that vitamin D can abrogate impairment caused by oxidative stress of nuclear translocation of the ligand bound glucocorticoid receptor [55].

This action of vitamin D to enhance antioxidant pathways and mitigate against oxidative stress likely contributes to its capacity to suppress IL-6 production by PM-stimulated HBECs. Consistent with this are our observed effects of sulforaphane in cell culture and the findings of Sienra-Monge *et al.* that antioxidant supplementation of children significantly decreases nasal lavage concentrations of IL-6 but not CXCL8 [56]. Oxidative stress has also been shown to contribute to corticosteroid resistance, a major feature of severe asthma [57], and we have previously shown that vitamin D exerts a steroid-sensitising action in severe asthma, restoring dexamethasone-induced IL-10 production by T cells [58].

However the capacity of vitamin D to suppress production of cytokines such as IL-6 is likely multifactorial since vitamin D also has the propensity to inhibit other signalling pathways, such as those involving nuclear factor kappa-B (NFκB) [21] and mitogen-activated protein kinase (MAPK) [59]. The differing effects of vitamin D on IL-6, CXCL8 and GM-CSF reveals the complex actions of vitamin D on the multiple intracellular signalling pathways in the induction of each cytokine. The suppression of IL-6 by vitamin D in our *in vitro* cultures is consistent with the previous finding of Codoner-Franch *et al.* that systemic concentrations of IL-6 in obese children were significantly higher in those with lower serum vitamin D concentrations [60].

A limitation of this research is that the detailed series of mechanistic experiments could only be conducted with epithelial cell cultures from a limited number of healthy and asthmatic donors. Furthermore the asthmatic donors were predominantly of mild severity and atopic; whether cells isolated from more severe and non-atopic asthmatics would respond similarly is an important consideration. The impact of asthma medications on these responses is also an important question but could not be analysed with this sample population. It is therefore reassuring that our findings are consistent with *in vivo* cytokine responses as discussed above [56] [60]. Furthermore our results are consistent with the epidemiological study of Rosser and colleagues that showed vitamin D insufficient children living close to major roads have an increased risk of severe asthma exacerbations [12]. Importantly the concentrations of vitamin D and particulate matter used in this research are representative of the concentrations evident in the real world. Li *et al.* have reconciled theoretical and experimental particulate airway deposition, allowing for increased deposition at airway bifurcation points with uneven air flow in disease, and shown PM_2.5_ deposition concentrations of 0.2–20 μg/cm^2^ are representative for real-world exposure [18] with higher deposition concentrations for total particulate matter. Circulating serum levels of 25-hydroxylated vitamin D are in the range 75nmol – 150nmol in sufficient individuals, and therefore an *in vitro* concentration of 100nmol 25(OH)D3 is representative of real-world vitamin D sufficiency. With respect to 1,25(OH)_2_D3 there is greater uncertainty as to *in vivo* tissue concentrations but the *in vitro* effects of 100nm 1,25(OH)_2_D3 on immunological markers mirror *ex vivo* correlates supporting this concentration being representative [19].

Although there are very few studies examining a beneficial role for vitamin D in alleviating adverse effects of pollution, there is a larger volume of evidence examining antioxidant supplementation as a strategy to reduce the harm to human health from air pollution. For example multiple studies have investigated the capacity of antioxidant supplements (such as vitamin C and vitamin E) to abrogate any deleterious effect of air pollution on lung function and many of these have shown significant benefit [61]. Similarly there is suggestive evidence that supplementation of vitamin C and E may improve athletic performance in ozone-exposed runners [62], and that B vitamin supplementation can alleviate particulate exposure associated cardiac autonomic dysfunction [63], amongst other outcome measures studied. The major difference between vitamin D and other antioxidant vitamins (and minerals) is that the major natural source for vitamin D in humans is through sunlight exposure not diet. Vitamin D insufficiency / deficiency are extremely common in the developed world [64] and a healthy diet alone rarely contains adequate vitamin D to achieve sufficient status. Although ultraviolet (UV) light supplementation might be an option [65], population-level oral vitamin D supplementation may be necessary to alleviate the epidemic of vitamin D deficiency and reduce harm from air pollution.

Our study provides mechanistic evidence to support larger trials of vitamin D supplementation to alleviate the harmful impact of air pollution, similar to those conducted with other antioxidant vitamins. Our conclusions are based on samples from a limited number of subjects and next need to be followed up in larger translational studies. In the first instance exposure chamber studies in vitamin D deficient individuals randomised to vitamin D supplementation / placebo prior to exposure would provide the opportunity to confirm whether *in vivo* vitamin D supplementation beneficially impacts on the pulmonary response to ambient pollutants. Clinical endpoints and biomarkers could be studied in a larger sample of participants including healthy controls, a broader range of asthmatic patients and patients with other respiratory diseases.

This research focused on the effects of vitamin D on pollution-induced pro-inflammatory cytokine production by human bronchial epithelial cells, through the upregulation of antioxidant defences. Transcriptomics was used to identify cytokines of interest, with the advantage of revealing differentially modulated cytokines in an unbiased manner. The array also identified multiple non-cytokine disease-associated genes regulated by PM-stimulation and vitamin D - such as *CLDN7, SERPINB1, COL1A1, SLPI* and *MMP9* - that were beyond the scope of this research. Many of these have been implicated in airways pathology and deserve future research.

## Conclusions

In summary, vitamin D decreased oxidative stress and the particulate matter-induced IL-6 response in HBEC cultures; with vitamin D pre-treatment suppressing the IL-6 response consistently in cultures from both healthy and asthmatic donors. Therefore vitamin D sufficiency is likely beneficial in protecting against pollution-induced inflammation in asthma, as well as in other pollution-associated diseases related to the induction of airway and systemic inflammation. Vitamin D supplementation has been shown to be safe and effective in preventing severe asthma exacerbations and acute respiratory tract infections when given to appropriate target groups [66] [67]. This research mechanistically supports a strategy of public health intervention to optimise the body’s own defences against pollution (for example supplementation with vitamin D [12] or other vitamins [63]) to reduce the burden of pollution-associated disease.

## Acknowledgements

The authors would like to acknowledge scientific advice pertaining to this research theme from Dr Nicholas Matthews and assistance in recruiting participants from Kheem Jones.

## Supporting Information Captions

**S1 Table. Taqman primer probesets**.

**S2 Table. Genes showing differential regulation by both NIST stimulation and vitamin D in the transcription microarray**.

Gene expression fold-changes in a transcription microarray of HBECs cultured for 24 hours with/without 50μg/ml NIST stimulation in the presence/absence of 100nM 1,25(OH)2D3. Short-list of genes showing both differential expression with NIST stimulation (fold change ≥ ± 1.4) and differential expression with vitamin D treatment in presence of NIST stimulation (fold change ≥ ± 1.4).

**S1 Fig: Cytokine production by HBEC cultures from healthy and asthmatic donors stimulated by NIST PM with / without 1,25(OH)_2_D3**.

Primary HBECs stimulated for 24 hours with 50μg/ml NIST in the presence/absence of 100nM 1,25(OH)_2_D3. VC; vehicle control for NIST. Healthy donor cultures: n=8 for IL-6, n=9 for CXCL8, n=8 for GM-CSF. Asthmatic donor cultures: n=7 for IL-6, n=8 for CXCL8, n=6 for GM-CSF. Friedman’s tests with Dunn’s multiple-comparisons tests. Statistical significance: *, p ≤ 0.05.

**S2 Fig: Induction of cytokine gene expression in HBEC cultures from healthy and asthmatic donors stimulated by NIST PM with / without 1,25(OH)_2_D3**.

(A) Induction of *IL6* and *CSF2* by 50μg/ml NIST PM stimulation in the presence / absence of 100nM 1,25(OH)_2_D3 in 4 hour and 24 hour cultures of HBECs from healthy and asthmatic donors. H, Healthy donor HBECs; A, Asthmatic donor HBEC cultures. Tie-bars show paired results from the same donor cultures. 4hr healthy, n=5; 4hr asthmatic, n=5; 24hr healthy, n=6-7; 24hr asthmatic, n=6.

(C) Induction of *IL6* and *CSF2* by NIST at 4 hours compared to suppression by 100nM 1,25(OH)_2_D3 at 24 hours of IL-6 and GM-CSF production respectively in NIST PM stimulated HBEC cultures. n=8 (4 healthy and 4 asthmatic).

**S3 Fig: Effect of NIST stimulation on expression of the vitamin D axis genes *CYP27B1* and *VDR*, and comparison of response to 25(OH)D3 and 1,25(OH)_2_D3**.

(A) Expression of *CYP27B1* mRNA in 24 hour HBEC cultures stimulated with 50μg/ml NIST PM with/without 100nM 1,25(OH)_2_D3 (n=9), or with 1μg/ml Poly(I:C) (paired t-test, n=6). Gene expression measured by qPCR relative to the house-keeping gene *18S*, and shown relative to the control condition. Statistical significance as follows: *, p ≤ 0.05; **, p ≤ 0.01; ***, p ≤ 0.001.

(B) Expression of *VDR* in 24 hour cultures stimulated with PM with/without 1,25(OH)_2_D3 (n=7), or with 1μg/ml Poly(I:C) (paired t-test, n=4).

(C) Percentage suppression of cytokine-production in 24 hour cultures of PM-stimulated HBECs upon concurrent addition of 1,25(OH)_2_D3 or 25(OH)D3. Cultures from healthy donors shown as open circles and those from asthmatic donors as filled squares. IL-6, n=7 healthy, 6 asthmatic; CXCL8, n=8 healthy, 7 asthmatic; GM-CSF, n=6-7 healthy, 6 asthmatic.

**S1 Data Appendix. Transcription microarray results after quantile normalisation and ANOVA analysis (full gene list)**.

